# Amygdalar Calcitonin Gene-Related Peptide Driven Effects of Cold Sensitivity Induced by Peripheral Neuropathy in Mice

**DOI:** 10.1101/2025.05.23.655813

**Authors:** Alexis D. Trail, Heather N. Allen, Tyler S. Nelson, James A. Widner, Lakeisha Lewter, See H. Tack, Benedict J. Kolber

**Author notes:** COMMUNICATING AUTHOR: Dr. Benedict Kolber University of Texas at Dallas 800 W. Campell Road Richardson, TX 75080.

## Abstract

The central nucleus of the amygdala (CeA) is a critical regulator of nociception, and its role in pain modulation depends on factors such as hemispheric location, neuropeptide release, and experimental model. Calcitonin gene-related peptide (CGRP) is a potent neuropeptide modulator within the CeA. Previous research has demonstrated its CeA nociceptive role in migraine, visceral, arthritic, and inflammatory pain murine models. The contribution of CeA CGRP to neuropathic pain is unclear. This study examined the effects of CGRP and its receptor antagonist, CGRP 8-37, in the CeA on mechanical and cold sensitivity in two mouse models of neuropathic pain: chemotherapy-induced peripheral neuropathy (CIPN) mediated by paclitaxel (PTX) and injury-induced neuropathy through the spared nerve injury (SNI) model. Mechanical and cold sensitivity were measured using the hindpaw von Frey and topical acetone drop assays, respectively. Neither CGRP nor CGRP 8-37 in the CeA had any significant effect on mechanical sensitivity in either neuropathic pain model. In the SNI-treated mice, CGRP infusion into either the left or right CeA reduced cold sensitivity in the left and right SNI-treated hindpaw, while CGRP 8-37 infusion into the left or right CeA increased cold sensitivity in the right SNI-treated hindpaw only. In PTX-treated mice, CGRP infusion into the left or right CeA decreased cold sensitivity of the contralateral paw only. These results suggest that CGRP in the CeA influences pain modulation in a complex manner that depends not only on the hemisphere and injury site, but also on the underlying cause of the neuropathic condition.

**PERSPECTIVE:** This article presents the anti-nociceptive properties of calcitonin gene-related peptide (CGRP) signaling within the central nucleus of the amygdala during neuropathic pain-like conditions in mice. This dataset can serve to guide novel drug development for treating chronic neuropathic pain conditions.

**HIGHLIGHTS:**

- Lateralization of the central nucleus of the amygdala (CeA) is pain-model dependent
- CGRP signaling within the CeA is correlated with decreased cold sensitivity
- Different neuropathic etiologies yield differences in CGRP responsiveness

## INTRODUCTION

Chronic neuropathic pain is caused by injury or dysfunction of the somatosensory nervous system that affects about 10% of adults.^1^ It may result from a range of underlying conditions, such as diabetes, autoimmune disorders, viral infections, cancer, chemotherapy, or mechanical damage to nerves.^2^ Treatment for chronic neuropathic pain varies, with the main treatment options being gabapentinoids, tricyclic antidepressants (TCA), serotonin–norepinephrine reuptake inhibitors (SNRI), topicals, botulinum toxin injections, or opioids.^3^ Although these options are sufficient for some, many patients do not experience adequate pain relief from these treatment options or must endure undesirable side effects^3^, demonstrating a clinical need to develop novel treatments for chronic neuropathic pain that are effective and safe.

Chronic pain is a complex, multi-dimensional experience involving sensory-discriminative, emotional, and cognitive components, all of which are shaped by dynamic brain circuitry. Given the high prevalence of affective comorbidities among individuals with chronic pain^4^, it is critical to study the neural mechanisms linking the sensory and emotional aspects of pain. One key region of interest for this link is the amygdala, a small, subcortical structure within the medial temporal lobe. The amygdala is known for its roles in aggression^5^, fear conditioning^6^, emotional regulation^7^, and pain^8^. Within the amygdala, the central nucleus (CeA) has emerged over the last 30 years as a critical node for nociceptive modulation. The CeA receives direct nociceptive input from the parabrachial nucleus (PBN) and plays a pivotal role in descending pain-modulatory circuits.^9–11^ Many previous studies have further explored the CeA’s overall involvement in the modulation of pain, including lesion^12–18^, optogenetic^19–21^, chemical inactivation^22–26^, and electrical^27,28^ or chemical^13,26,29,30^ activation studies indicating the CeA is pro-nociceptive^12,13,17,23,24,29,30^, anti-nociceptive^14–16,18,21,22,25,27,28^, or situationally both^19,20,26^.

Interestingly, the CeA’s role in pain modulation is situationally dependent on hemispheric location^8,20,24,25,31,32^. Current studies demonstrate that the left or right CeA can be pro-nociceptive, anti-nociceptive, or have no effect on nociception, depending on factors such as the contralateral/ipsilateral orientation of injury to the CeA, pain model implemented, nociceptive phenotype measured, or neuropeptide manipulated^8^. The CeA’s neuropeptide population is a particularly interesting source of functional variance in nociception^33^. Previous studies have explored the role of specific neuropeptide signaling within the CeA in pain modulation, such as corticotropin releasing factor (CRF)^32,34–43^, somatostatin (SST)^19,26^, pituitary adenylate cyclase activating polypeptide (PACAP)^44,45^, and to this study’s interest, calcitonin gene-related peptide (CGRP)^30,31,46–51^. CGRP is a vasodilator expressed throughout multiple regions and organ systems within the body, including the cardiovascular system^52^, enteric system^53^, and the nervous system^54^. Within the CNS, CGRP is heavily expressed in regions associated with stress and pain processing, including the periaqueductal grey, PBN, locus coeruleus, and the amygdala.^55^ CGRP is a promising target for the central modulation of pain, as previous studies have discovered its direct involvement in nociception, particular in CGRP originating from the PBN.^56–58^ CGRP-containing afferent neurons of the PBN bind to CeA neurons expressing CGRP receptors, establishing CGRP as a part of the spino-parabrachial-amygdaloid pain pathway.^59,60^ However, the past evaluation of amygdalar CGRP on nociception has been limited to naïve animals or migraine, visceral, arthritic, inflammatory, and spinal nerve ligation (SNL) neuropathic pain modeled animals, of which the results are varied between pain-like phenotypes.^30,31,46–51^ These data include those showing that CGRP can be either pro-nociceptive^30,46–48,50,51^, anti-nociceptive^49^, or situationally both^31^ in the CeA.

A current gap in understanding the role of CGRP in the CeA on modulation of neuropathic nociception is the potential factor of hemispheric lateralization. Although a previous study evaluated CGRP signaling in the right CeA of SNL neuropathic pain modeled animals^51^, the potential effect(s) of the left CeA was not tested. Our study aimed to fill that gap while additionally considering potential effects mediated by different neuropathic pain etiologies, including the injury-based spared nerve injury (SNI) model and paclitaxel (PTX)-mediated chemotherapy-induced peripheral neuropathy (CIPN) model. Given the etiological differences of the SNI and CIPN neuropathic conditions, we hypothesized there may be a difference in the effect of amygdalar CGRP on nocifensive responses in these models with potential for hemispheric lateralization to modulate these differences..

## METHODS

### Animals

A total of 32 female and 44 male 9–12-week-old wildtype C57BL/6J mice were used across all experiments. All animals were single-housed for at least 7 days on a 12-hour light/dark cycle and provided rodent chow and water *ad libitum* after surgery and prior to behavioral testing. Animals were maintained, and experiments were conducted in accordance with the Institutional Animal Care and Use Committees at the University of Texas at Dallas (Richardson, TX; protocol 20-04) and Duquesne University (Pittsburgh PA; protocol 2006-03). Animals were bred in-house from C57BL/6J Jackson breeders refreshed after two in-house generations of breeding. All efforts were made to minimize animal suffering and to reduce the number of animals used. Experimenters were blinded to drug treatment and hemispheric side of injection, but not sex, until all data were analyzed.

### Surgical procedure for cannula implantation

The mice were briefly anesthetized in an isoflurane induction chamber (3-5%) and transferred to a stereotaxic frame with an isoflurane and oxygen induction nose-cone (3% via SomnoSuite). A 7.97 mm (0.2 mm diameter) stainless steel cannula (Microgroup, Inc.) was implanted above the right and/or left central nucleus of the amygdala (CeA) (bregma: AP:-1.45 mm, midline: ±3.00 mm, DV (from skull): –4.2mm) while the mice were continuously anesthetized with 3% isoflurane with oxygen. Two bone screws were attached to the skull above bregma and lambda, and dental cement was used to securely fix the cannula and bone screws to the skull. The back of the scalp was sutured together over the dental cement to prevent skin growth under the skull cap. Post-surgical recovery treatment included the application of 2.5%-2.5% lidocaine/prilocaine cream (Alembic Pharmaceuticals NDC# 62332058204) to the scalp and a single 3.25 mg/kg intraperitoneal injection of buprenorphine HCl (Covetrus CAT# 059122). The mice were given >1 week of recovery prior to behavioral testing. Correct targeting was determined at the end of the experiment. The brain was extracted, and the physical location of the cannula was evaluated by cutting down the cannula track and evaluating the tip of the cannula site. Off-target surgeries were not included in the analysis. A total of 3 animals (2 male,1 female) were excluded from analysis due to an off-target cannula.

### Drug treatment preparation and administration

On pharmaco-behavioral testing days, mice received a 1 uL injection of 1x aCSF (artificial cerebrospinal fluid), 100 uM CGRP (GenScript RP11095, lot E5669022E) reconstituted in 1x aCSF, or 100 uM CGRP 8-37 (CGRP receptor antagonist) (GenScript RP11090, lot E286662K) reconstituted in 1x aCSF. The final drug dosage for CGRP and CGRP 8-37 was 100 pmol per injection. Dosing of drugs was based on a previous publication modulating CGRP in the context of bladder pain^31^. 1x aCSF was made fresh within 24 hours of preparing or administering drug. 1x aCSF was made by diluting 10x aCSF with ultrafiltered MilliQ water (1:10). 10x aCSF was prepared using the following reagent concentrations dissolved in ultrafiltered MilliQ water: 25 mM KCl, 12.5 mM NaPO4·H2O, 1.25 M NaCl, 25 mM NaHCO3, 20 mM CaCl2, 13 mM MgCl2, and 100 mM Dextrose. 10x aCSF and 1x aCSF were filtered after combining reagents and brought to a pH of 7.4. In a cross over design, all mice received each drug treatment in the order defined in **Table 1**. The drug treatment schedule was randomized for each cohort and behavioral testing was completed blinded to treatment. The experimenter conducting behavioral tests for the SNI experiment was additionally blinded to the hemispheric site of injection since these were bilateral cannulations. Prior to intra-amygdalar injection via cannula, mice were anesthetized in an isoflurane induction chamber, then subsequently moved to an induction nose-cone where they received 2.0-2.5% isoflurane vaporized in 100% oxygen for the duration of the injection lasting 10 minutes. Drug was administered at a rate of 0.1 uL/min using a 32-gauge injection cannula that extended 0.1 mm below the end of the cannula, coupled to a Hamilton syringe fixed to a syringe pump via flexible plastic tubing through which drug flowed.

**Table 1.**
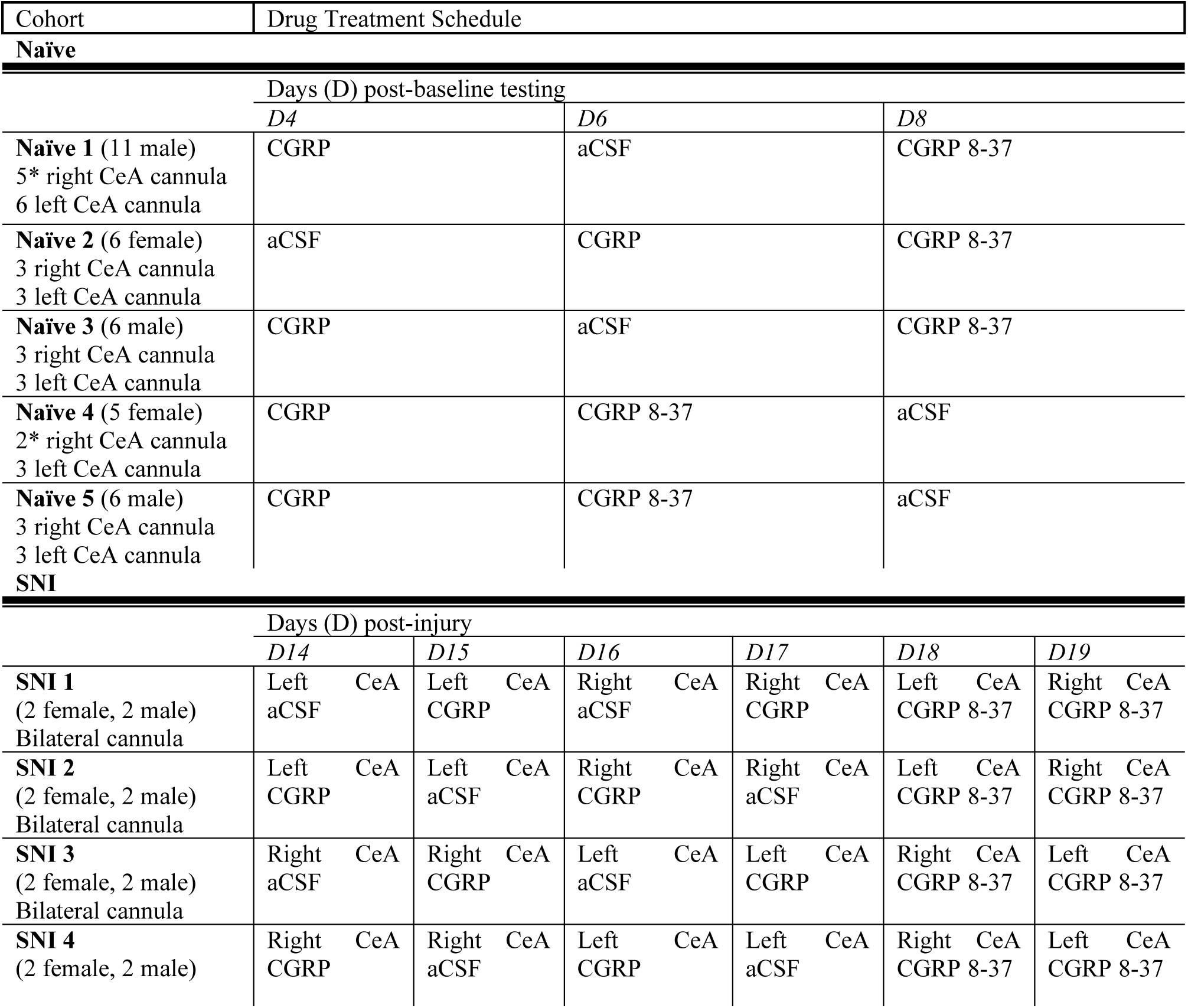
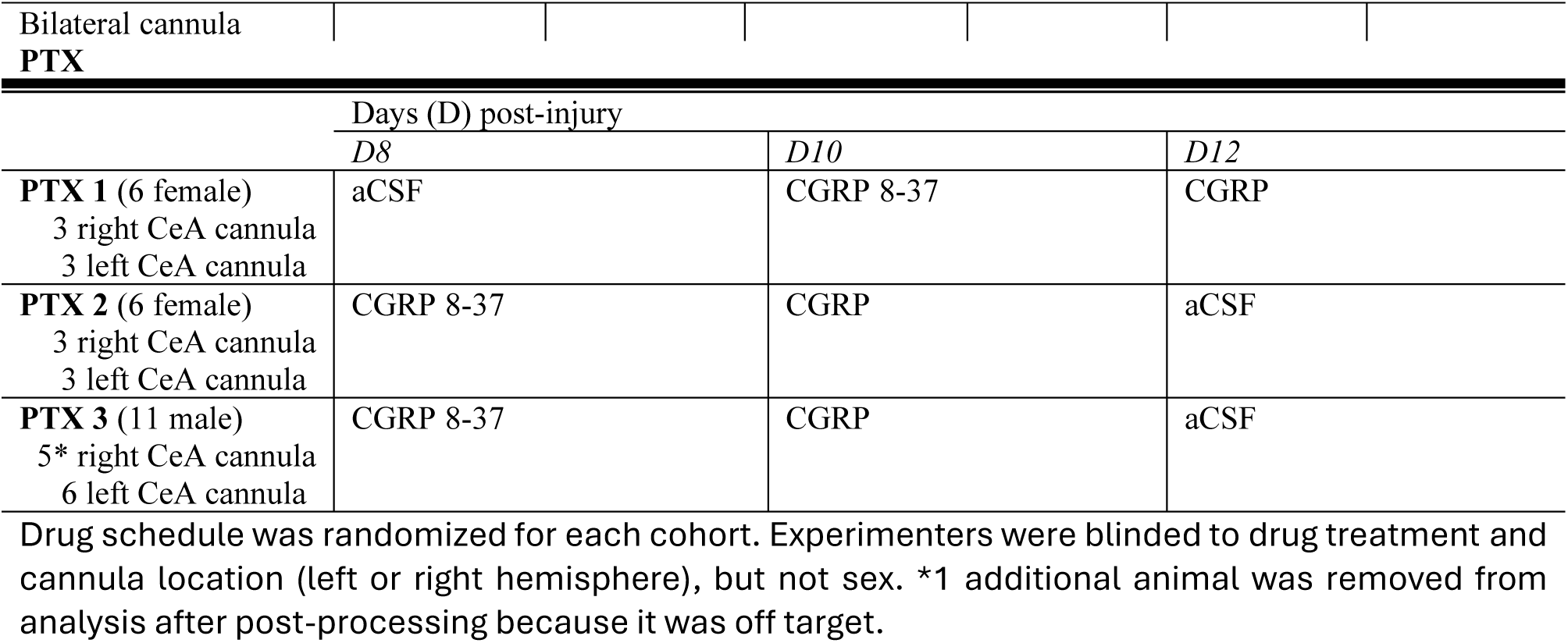
Drug treatment schedule.

### Chemotherapy-induced peripheral neuropathy (CIPN)

PTX injections were used to induce CIPN in unilaterally cannulated mice. PTX stock solution was made by combining solid PTX (Selleckchem S1150, lot S115012) with a 1:1 kolliphor and 100% ethanol solution to create a 6 mg/mL PTX solution. On PTX injection days, the stock solution was further diluted with sterile saline to a 0.6 mg/mL concentration of PTX. Mice were given a 4 mg/kg dose of the diluted PTX solution injected intraperitoneally (i.p.) every other day for 7 days for a total of 4 doses, resulting in a cumulative dose of 16 mg/kg per mouse.^61^ Experimental behavioral testing began 24 hours after the final dose which would be considered Day 8 (D8). Behavioral testing was completed every other day for 6 days following the end of PTX treatment.

### Spared nerve injury (SNI)

In bilaterally-cannulated mice, the SNI protocol was used to induce a one-sided peripheral injury that models neuropathic pain as previously described.^62^ Mice were anesthetized in an induction chamber (5% isoflurane) and subsequently placed on a nose-cone where they received 2% isoflurane vaporized in 100% oxygen. The tibial and common peroneal branches of the sciatic nerve were ligated and cut while the sural branch was left unmanipulated.^63^ SNI was performed on the final day of baseline testing, 1 week after cannulation. SNI was completed on the left or right sciatic nerve. 14 days after SNI, pharmaco-behavioral testing was conducted over 6 days. The injected drug for each animal was randomized on each day of testing and alternated between the left and right CeA as described in **Table 1**.

### Mechanical and cold sensitivity testing

Mechanical and cold sensitivity was assessed using the up-down method with von Frey filaments applied to the hindpaw^64^ and topical acetone application to the plantar surface^65^, respectively. Mechanical sensitivity was quantified by calculating the 50% withdrawal threshold (in grams) for both hindpaws, and cold sensitivity was quantified using the total duration of paw withdrawal (seconds) of both hindpaws within 60 seconds after acetone application. The following von Frey filaments (TouchTest) were used: 0.02g (0.19mN), 0.04g (0.39mN), 0.08g (0.78mN), 0.16g (1.5mN), 0.32g (3.1mN), 0.64g (6.3mN), 1.28g (12.6mN), 2.56g (25.1mN). Baseline data for these assays was acquired 1 week after cannula implantation when the mice were fully recovered from cannula surgery. For all behavioral testing, mice were allowed to habituate in the testing environment for ≥ 1 hour prior to testing. The experimenter was present in the room for the final 30 minutes of habituation. Female and male mice were tested separately. On pharmaco-behavioral testing days, baseline behavioral data was conducted after habituation and prior to drug administration. Drugs were administered under light anesthesia as described above and mice were returned to the testing apparatus to recover. Mice were not anesthetized immediately prior to pre-injury baseline and pre-drug baseline testing because there was no intra-cannula drug administration. Experimental behavioral data was collected 30-45 minutes after drug infusion, a time window previously identified as peak of effects for intra-amygdalar CGRP and its receptor antagonist CGRP 8-37.^31^. The testing order of the drugs was randomized and the experimenter was blinded to treatment. The experimenter conducting behavioral tests for the SNI experiment was additionally blinded to the hemispheric site of injection. Previous data shows that CGRP drug administration order has no effect on pharmaco-behavioral data^31^. The experimenter was not blinded to the sex of the mice.

### Statistics and data analysis

All data analysis was conducted blinded to treatment. The experimenter conducting the data analysis for the SNI cohort was additionally blinded to the hemispheric site of injection. Male and female data were not analyzed separately. Behavioral data was analyzed on GraphPad Prism (version 10.3.0) using ordinary one-way ANOVA or repeated measures two-way ANOVA followed by Šidák’s or Dunnett’s post-hoc multiple comparisons test as recommended by the software. Consistent with the majority of the pain literature, von Frey mechanical sensitivity data were analyzed using parametric statistics. The mechanical sensitivity datasets for the naïve, SNI, and PTX treated cohorts were excluded from the analysis in **Figure 4**, as there were no significant differences between pre-injection (i.e., baseline) and post-injection values (i.e., 50% withdrawal thresholds) for any of the three experiments. Statistical significance was determined at the level of P<0.05. Asterisks denoting P values correspond to: *P<0.05, **P<0.01, ***P<0.001, ****P<0.0001. All data are shown as mean +/-the standard error of the mean (SEM). Detailed information regarding statistical tests for each figure can be found in **Supplemental Table S1**.

## RESULTS

### Exogenous amygdalar CGRP is associated with an anti-nociceptive phenotype in the right hindpaw of naïve mice

The first experiment in this study aimed to evaluate the impact of exogenous CGRP or CGRP inhibition in the left or right CeA of uninjured (i.e., naïve) mice (**Figure 1A**). In the left CeA, exogenous CGRP had no effect on the mechanical sensitivity of either hindpaw (**Figure 1B, 1C, 1D, 1E**). Blockade of CGRP receptor activity in the left CeA, via infusion of peptide antagonist CGRP 8-37, resulted in a lower 50% withdrawal threshold for the right hindpaw (**Figure 1C**) compared to the pre-injection baseline. Interestingly, this pro-nociceptive effect was not observed for the left hindpaw of the same cohort of mice (**Figure 1B**). In the acetone test, neither CGRP nor CGRP 8-37 had an effect on the cold sensitivity of either hindpaw in naïve mice (**Figure 1D, 1E**).

**Figure 1.**
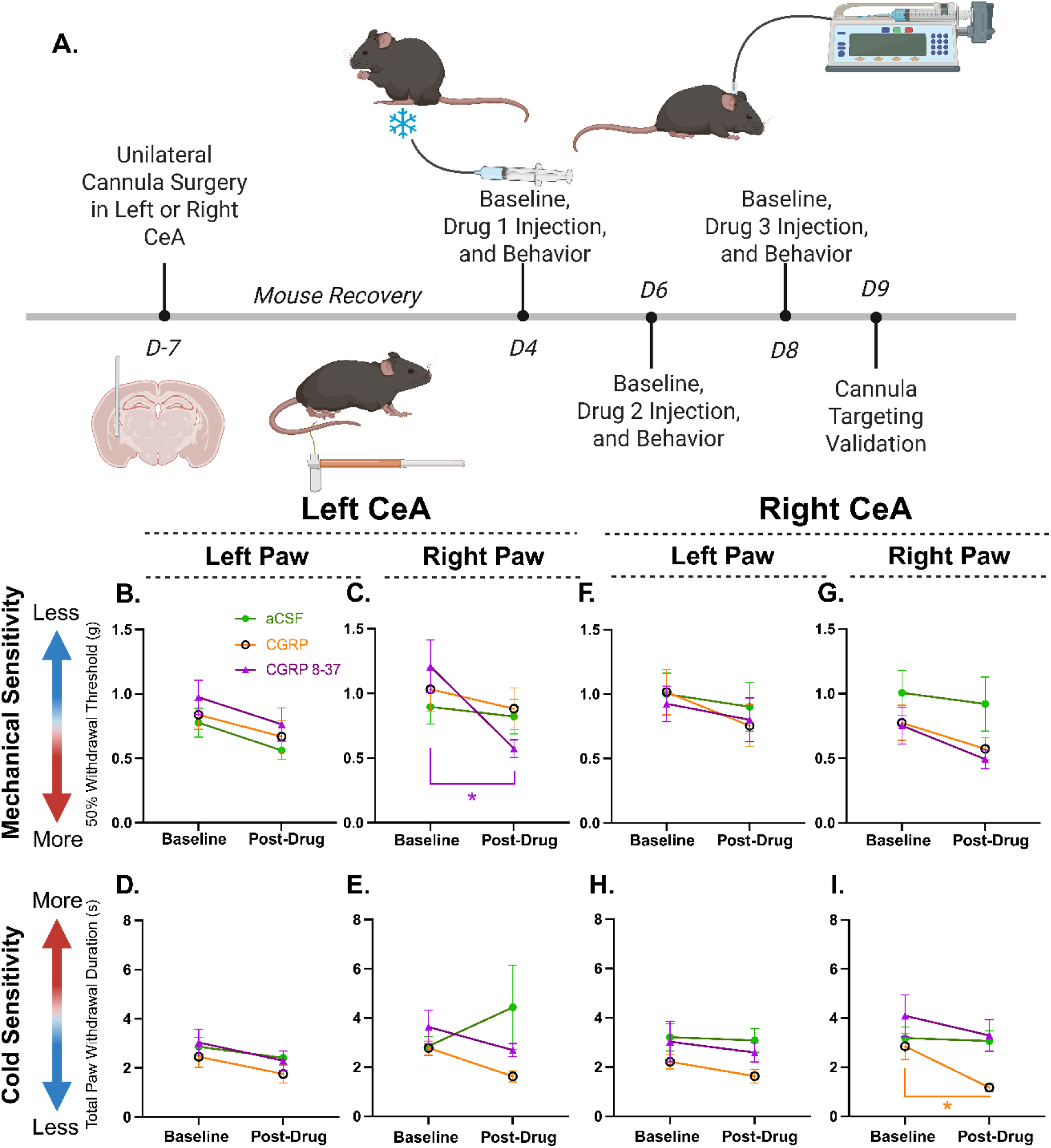
Injecting CGRP into the right CeA decreased cold sensitivity while CGRP 8-37 in the left CeA increased mechanical sensitivity in the right paw of naïve mice. **A)** Schematic representing the experimental timeline of intra-amygdalar cannula implantation, drug injections, and behavioral data collection of naïve mice. **B)** The 50% withdrawal threshold of the left (n=18) and **C)** right hindpaw (n=18) before and 30 minutes after injection of aCSF, CGRP, or CGRP 8-37 into the left CeA of naïve mice. **D)** The total paw withdrawal duration of the left (n=18) and **E)** right hindpaw (n=18) before and 30 minutes after injection of aCSF, CGRP, or CGRP 8-37 into the left CeA of naïve mice. **F)** The 50% withdrawal threshold of the left (n=16) and **G)** right hindpaw (n=16) before and 30 minutes after injection of aCSF, CGRP, or CGRP 8-37 into the right CeA of naïve mice. **H)** The total paw withdrawal duration of the left (n=16) and **I)** right hindpaw (n=16) before and 30 minutes after injection of aCSF, CGRP, or CGRP 8-37 into the right CeA of naïve mice. Data is represented as the mean with error bars representing the ± SEM. Statistical significance was determined using repeated measures two-way ANOVA with Šidák’s multiple comparisons test (* p<0.05, ** p<0.01, *** p<0.001, **** p<0.0001).

In the right CeA, neither CGRP nor CGRP 8-37 altered the mechanical sensitivity of either hindpaw (**Figure 1F, 1G).** In the acetone test, exogenous CGRP into the right CeA had an anti-nociceptive effect in the right hindpaw, as evidenced by a reduction of the total paw withdrawal duration when compared to the baseline test **(Figure 1I)**. However, there was no significant effect seen in the left hindpaw of the same cohort of mice (**Figure 1H**). Injection of CGRP 8-37 had no effect on cold sensitivity of either hindpaw **(Figure 1H, 1I)**.

### In the SNI model of neuropathic pain, exogenous amygdalar CGRP is associated with anti-nociception in response to cold but not mechanical stimuli

Next, this study aimed to evaluate the impact of exogenous CeA CGRP manipulation on pain-like behavior after SNI to the left or right sciatic nerve (**Figure 2A**). SNI produced a neuropathic pain-like phenotype in female and male mice, characterized by a significant increase in mechanical (**Figure 2B**) and cold (**Figure 2C**) sensitivity. In the left CeA, neither CGRP nor CGRP 8-37 altered the mechanical sensitivity of either SNI-treated hindpaw (**Figure 2D, 2E)**. In the acetone test, exogenous CGRP into the right CeA had an anti-nociceptive effect in the left and right SNI-treated hindpaw, as evidenced by a reduction of the total paw withdrawal duration when compared to the baseline test **(Figure 2F, 2G)**. Additionally, injection of CGRP 8-37 into the left CeA resulted in an increase in cold sensitivity for the right SNI-treated hindpaw only **(Figure 2F)**.

**Figure 2.**
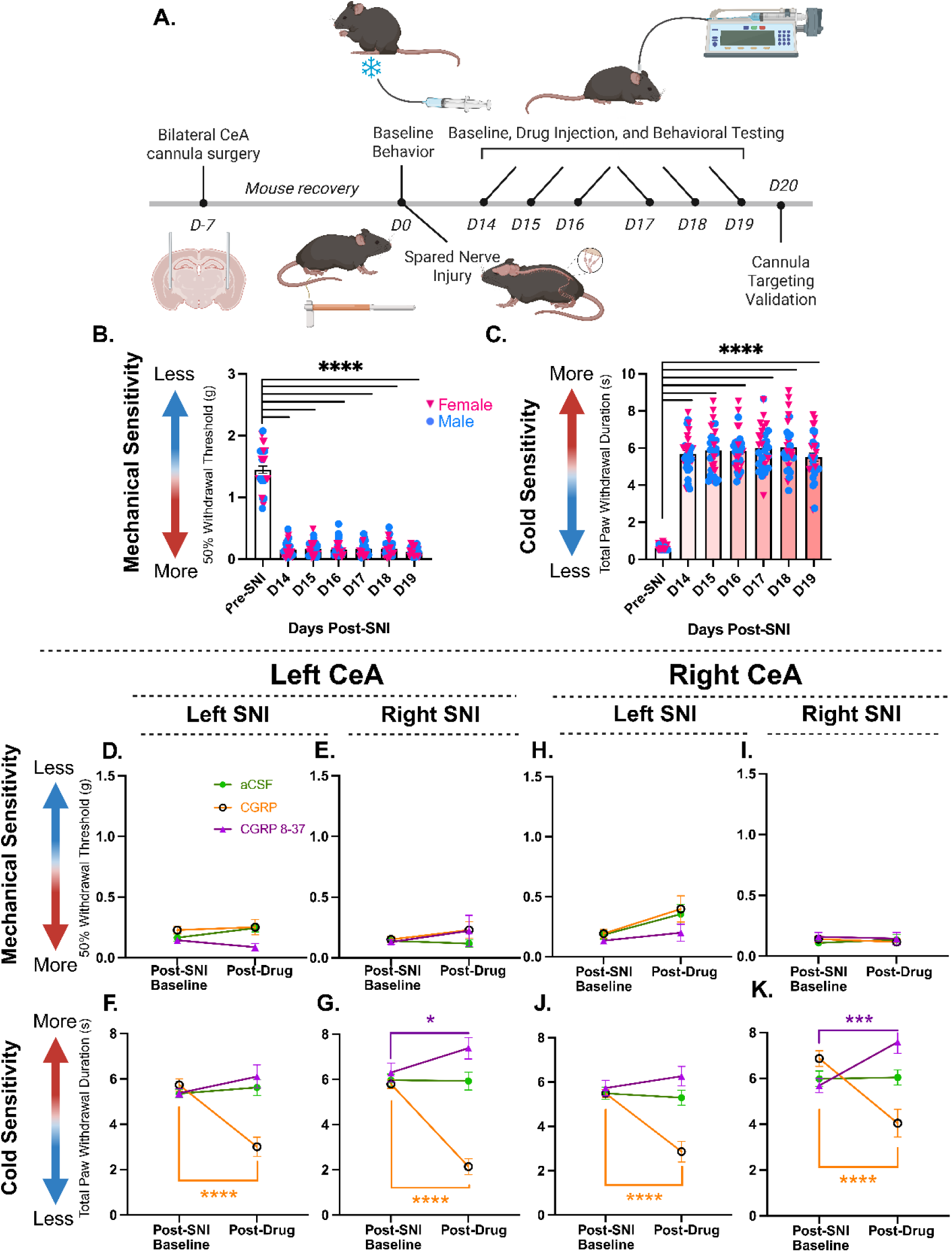
Injecting CGRP into the CeA decreased cold sensitivity in SNI-treated mice. CGRP 8-37 injected in the either the left or right CeA increased cold sensitivity only in the right SNI-injured paw. **A)** Schematic representing the experimental timeline of intra-amygdalar cannula implantation, drug injections, and behavioral data collection of SNI-treated mice. **B)** The average 50% withdrawal threshold and **C)** average total paw withdrawal duration of female (n=16) and male (n=16) mice pre– and post-SNI treatment. **D)** The 50% withdrawal threshold of the left (n=16) and **E)** right SNI-affected paw (n=16) before and 30 minutes after injection of aCSF, CGRP, or CGRP 8-37 into the left CeA of mice. **F)** The total paw withdrawal duration of the left (n=16) and **G)** right SNI-affected paw (n=16) before and 30 minutes after injection of aCSF, CGRP, or CGRP 8-37 into the left CeA of mice. **H)** The 50% withdrawal threshold of the left (n=16) and **I)** right SNI-affected paw (n=16) before and 30 minutes after injection of aCSF, CGRP, or CGRP 8-37 into the right CeA of mice. **J)** The total paw withdrawal duration of the left (n=16) and **K)** right SNI-affected paw (n=16) before and 30 minutes after injection of aCSF, CGRP, or CGRP 8-37 into the right CeA of mice. Data is represented as the mean with error bars representing the ± SEM. Statistical significance was determined using ordinary one-way ANOVA with Dunnett’s multiple comparisons test for graphs B-C and repeated measures two-way ANOVA with Šidák’s multiple comparisons test for graphs D-K (* p<0.05, ** p<0.01, *** p<0.001, **** p<0.0001).

In the right CeA, neither CGRP nor CGRP 8-37 altered the mechanical sensitivity of either SNI-treated hindpaw (**Figure 2H, 2I)**. However, exogenous application of CGRP in the right CeA resulted in a decrease in cold sensitivity of the left and right SNI-treated hindpaw, as evidenced by a reduction of the total paw withdrawal duration when compared to the baseline acetone test (**Figure 2J, 2K)**. Injection of CGRP 8-37 into the right CeA resulted in an increase in cold sensitivity for the right SNI-treated hindpaw only (**Figure 2K)**.

### In the PTX CIPN model, exogenous amygdalar CGRP is associated with anti-nociception in response to cold stimuli, but not mechanical stimuli, of the contralateral hindpaw

Finally, to evaluate whether the impact of exogenous CeA CGRP changed in the context of the etiology of the neuropathic model, a model of chemotherapy-induced peripheral neuropathy (CIPN) was used (**Figure 3A**). The chemotherapy agent PTX (i.p. 16mg/kg) produced a neuropathic pain-like behavioral phenotype in female and male mice, characterized by a statistically significant increase in mechanical (**Figure 3B**) and cold (**Figure 3C**) sensitivity. In the left CeA, neither CGRP nor CGRP 8-37 altered the mechanical sensitivity of either hindpaw (**Figure 3D, 3E)**. In the acetone test, CGRP into the left CeA had an anti-nociceptive effect in the right hindpaw, as evidenced by a reduction of the total paw withdrawal duration when compared to the baseline test **(Figure 3G)**. However, there was no significant effect seen in the left hindpaw of the same cohort of mice (**Figure 3F**).

**Figure 3.**
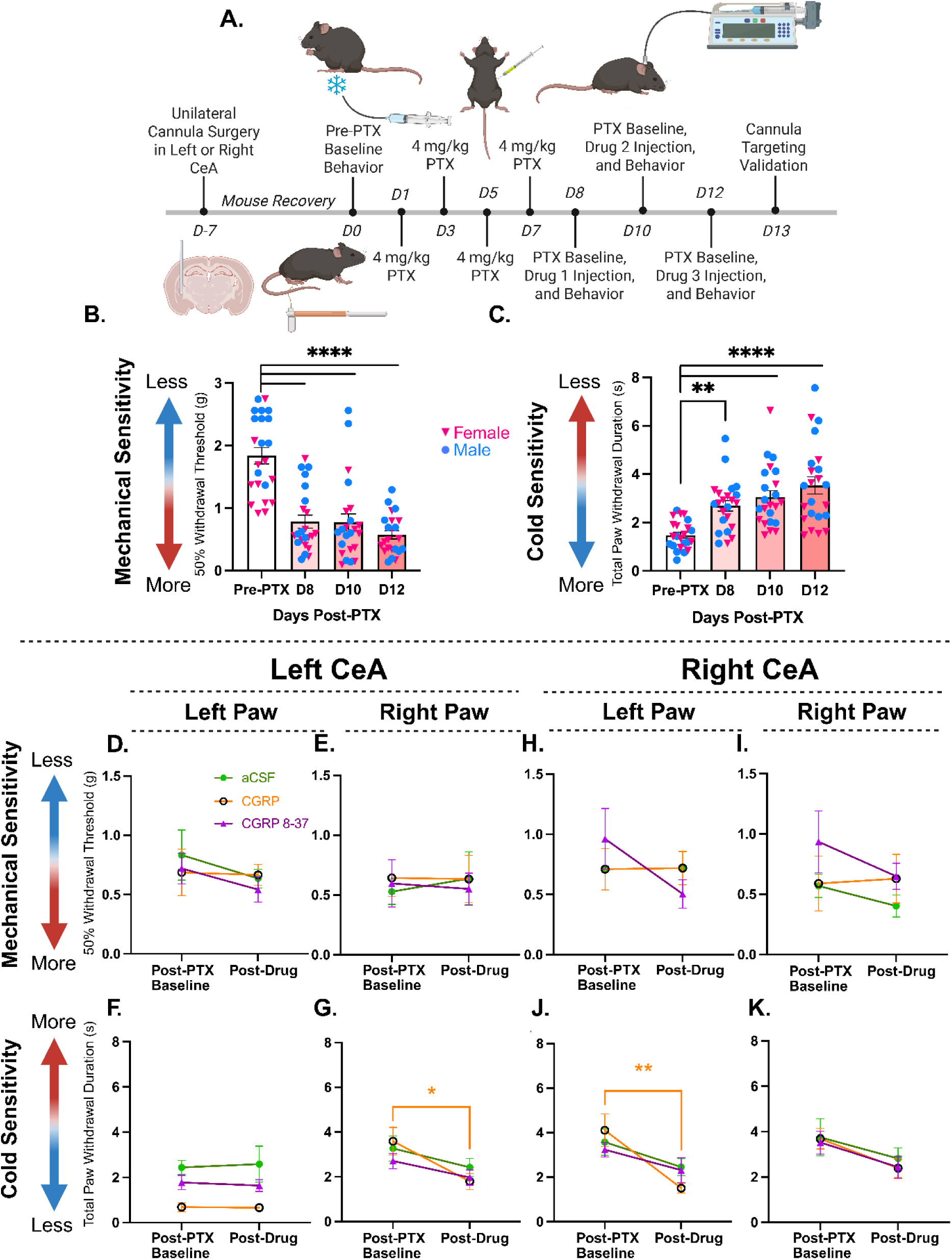
Injecting CGRP into the CeA decreased cold sensitivity in the contralateral paw of PTX-treated mice. **A)** Schematic representing the experimental timeline of intra-amygdalar cannula implantation, drug injections, and behavioral data collection of PTX-treated mice. **B)** The average 50% withdrawal threshold and **C)** average total paw withdrawal duration for female (n=12) and male (n=11) mice pre– and post-PTX treatment. **D)** The 50% withdrawal threshold of the left (n=12) and **E)** right hindpaw (n=12) before and 30 minutes after injection of aCSF, CGRP, or CGRP 8-37 into the left CeA of PTX-treated mice. **F)** The total paw withdrawal duration of the left (n=12) and **G)** right hindpaw (n=12) before and 30 minutes after injection of aCSF, CGRP, or CGRP 8-37 into the left CeA of PTX-treated mice. **H)** The 50% withdrawal threshold of the left (n=11) and **I)** right hindpaw (n=11) before and 30 minutes after injection of aCSF, CGRP, or CGRP 8-37 into the right CeA of PTX-treated mice. **J)** The total paw withdrawal duration of the left (n=11) and **K)** right hindpaw (n=11) before and 30 minutes after injection of aCSF, CGRP, or CGRP 8-37 into the right CeA of PTX-treated mice. Data is represented as the mean with error bars representing the ± SEM. Statistical significance was determined using ordinary one-way ANOVA with Dunnett’s multiple comparisons test for graphs B-C and repeated measures two-way ANOVA with Šidák’s multiple comparisons test for graphs D-K (* p<0.05, ** p<0.01, *** p<0.001, **** p<0.0001

**Figure 4.**
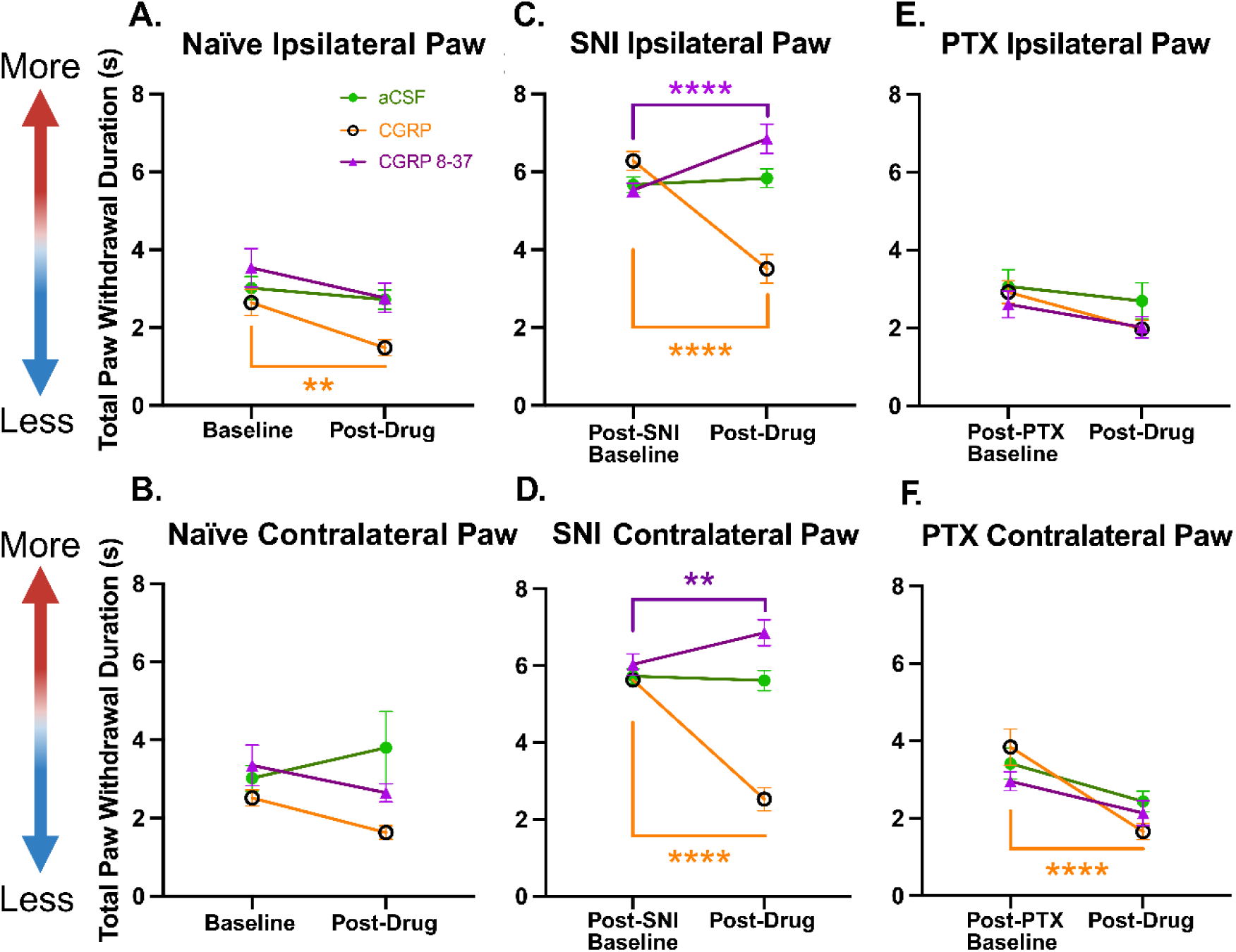
Injecting CGRP into the CeA decreased cold sensitivity in the ipsilateral paw of naïve mice and contralateral hindpaw of PTX-treated mice. CGRP in the CeA of SNI-treated mice decreased cold sensitivity while CGRP 8-37 increased cold sensitivity in the bilateral hindpaws. **A)** The total paw withdrawal duration of the ipsilateral (n=34) and **B)** contralateral hindpaw (n=34) to CeA cannula before and 30 minutes after injection of aCSF, CGRP, and CGRP 8-37 in naïve mice. **C)** The total paw withdrawal duration of the ipsilateral (n=32) and **D)** contralateral hindpaw (n=32) to CeA cannula before and 30 minutes after injection of aCSF, CGRP, and CGRP 8-37 in SNI-treated mice. **E)** The total paw withdrawal duration of the ipsilateral (n=23) and **F)** contralateral hindpaw (n=23) to CeA cannula before and 30 minutes after injection of aCSF, CGRP, and CGRP 8-37 in PTX-treated mice. Data is represented as the mean with error bars representing the ± SEM. Statistical significance was determined using repeated measures two-way ANOVA with **Š**idák’s multiple comparisons test (* p<0.05, ** p<0.01, *** p<0.001, **** p<0.0001).

In the right CeA, neither CGRP nor CGRP 8-37 altered the mechanical sensitivity of either hindpaw (**Figure 3H, 3I)**. In the acetone test, CGRP into the right CeA had an anti-nociceptive effect in the left hindpaw, as evidenced by a reduction of the total paw withdrawal duration when compared to the baseline test **(Figure 3J)**. However, there was no significant effect seen in the right hindpaw of the same cohort of mice (**Figure 3K**).

### Exogenous amygdalar CGRP modulates nociception in the ipsilateral, contralateral, or bilateral hindpaw(s) differentially based on pain state and etiology

To evaluate whether amygdalar CGRP’s nociceptive modulatory function is related to an ipsilateral or contralateral effect irrespective of “left” and “right”, the acetone test datasets from **Figures 1-3** were re-analyzed and graphed to combine the ipsilateral and contralateral CeA-hindpaw pathway effects, irrespective of side of brain. The mechanical sensitivity datasets were excluded from this analysis due to a lack of significant effects for any of the pain states. Analysis of the ipsilateral and/or contralateral effects on cold sensitivity, however, revealed an interesting trend that is differentially dependent on the modeled pain state. In naïve mice, intra-amygdalar injection of CGRP resulted in a significant reduction in cold sensitivity in the paw ipsilateral to the manipulated CeA (**Figure 4A**) and no significant effect in the paw contralateral to the manipulated CeA (**Figure 4B**). CGRP 8-37 within either CeA of naïve mice had no significant effect. In SNI-treated mice, intra-amygdalar injection of CGRP resulted in a significant reduction in cold sensitivity in the hindpaw bilaterally (**Figure 4E, 4F**). Additionally, CGRP 8-37 injection into the CeA resulted in a significant increase in cold sensitivity in the hindpaw bilaterally (**Figure 4E, 4F**), with a stronger pro-nociceptive effect seen in the paw ipsilateral to the manipulated CeA (**Figure 4E**). In PTX-treated mice, intra-amygdalar injection of CGRP resulted in a significant reduction in cold sensitivity in the hindpaw contralateral (**Figure 4D**), but not ipsilateral (**Figure 4C**), to the manipulated CeA while CGRP 8-37 had no significant effect. Detailed information regarding statistical analysis for each graph is provided in **Supplementary Table S1**.

## DISCUSSION

The goal of this investigation was to gain further insight into the role of amygdalar CGRP signaling in nociception by determining the dynamics of mechanical and cold modulation mediated by CGRP in the left or right CeA in a lesion-induced (SNI) and drug-induced (PTX) model of neuropathic pain compared to naïve conditions. The first aim was to determine the effects of exogenous CGRP manipulation in an uninjured (naïve) state to later compare to the neuropathic conditions. In naïve mice, CGRP in the right CeA decreased cold sensitivity while CGRP 8-37 in the left CeA increased mechanical sensitivity. Next, we found that regardless of the neuropathic pain-state tested, CGRP, in either the left or right CeA, has an anti-nociceptive effect and CGRP 8-37 has a pro-nociceptive effect when observing cold sensitivity. Interestingly, however, it appears that exogenous amygdalar CGRP signaling in the neuropathic pain models do not have a substantial effect on mechanical hypersensitivity in the present study, contrary to a previous report that found blockade of CGRP signaling via CGRP 8-37 to decrease mechanical sensitivity in a spared nerve ligation model of neuropathic pain in rats^51^.

These findings indicate that the relationship between CGRP and nociceptive modulation is more complicated than initially postulated. Prior to this investigation, the nociceptive nature of CGRP within the CeA was limited to naïve animals^47,49^ or rodent models of migraine^46^, visceral^31^, arthritic^48^, inflammatory^30,50^, and a SNL model of neuropathic pain^51^ summarized in **Table 2**. Combined with the data presented in the present manuscript it is not possible to conclude whether CGRP can be considered an anti-nociceptive, pro-nociceptive, or non-nociceptive neuropeptide within the CeA. The nociceptive function of CGRP varies heavily based on the type of pain model evaluated and the hemispheric orientation of the CeA, which was understudied in previous investigations.

**Table 2.**
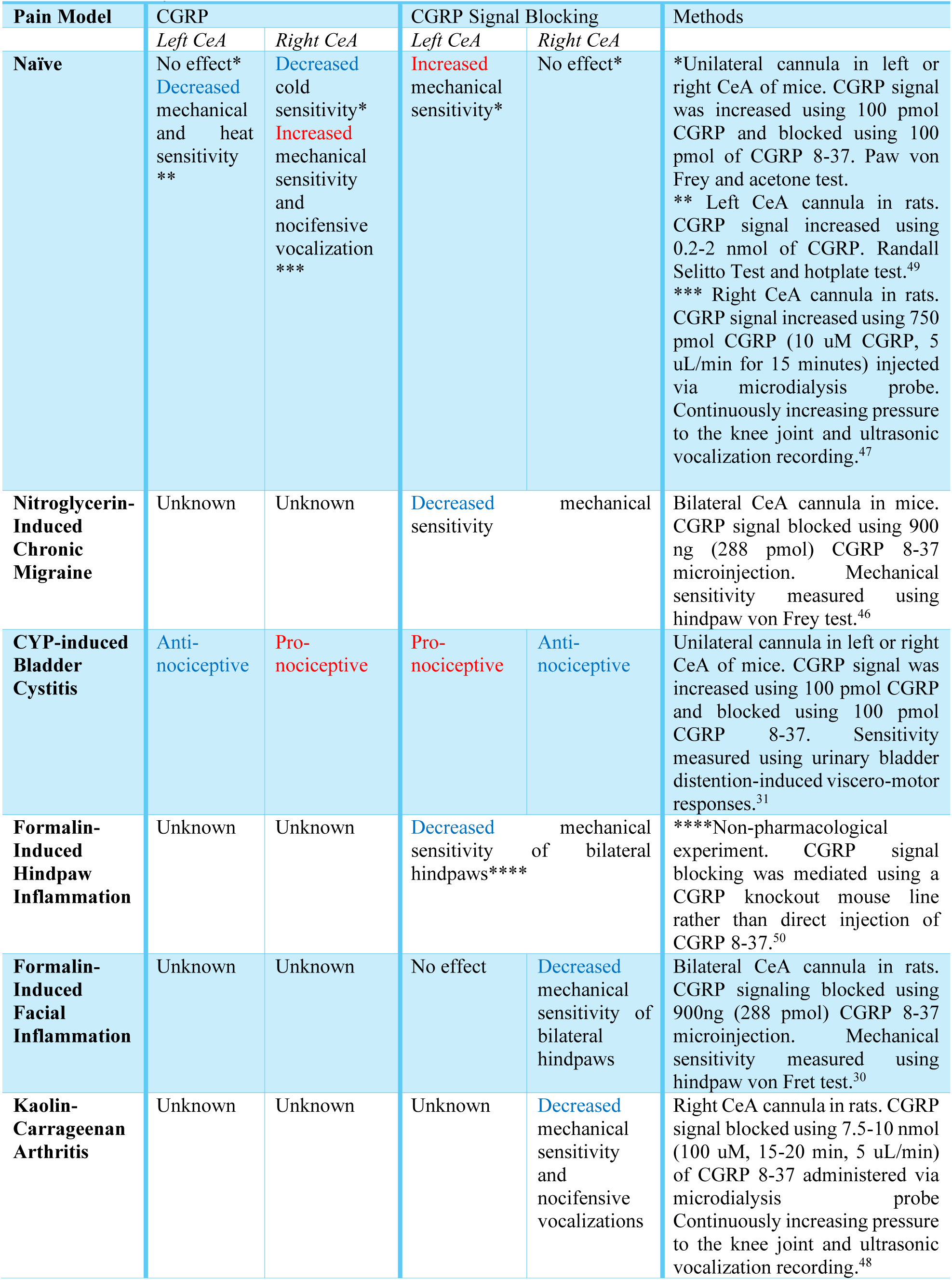
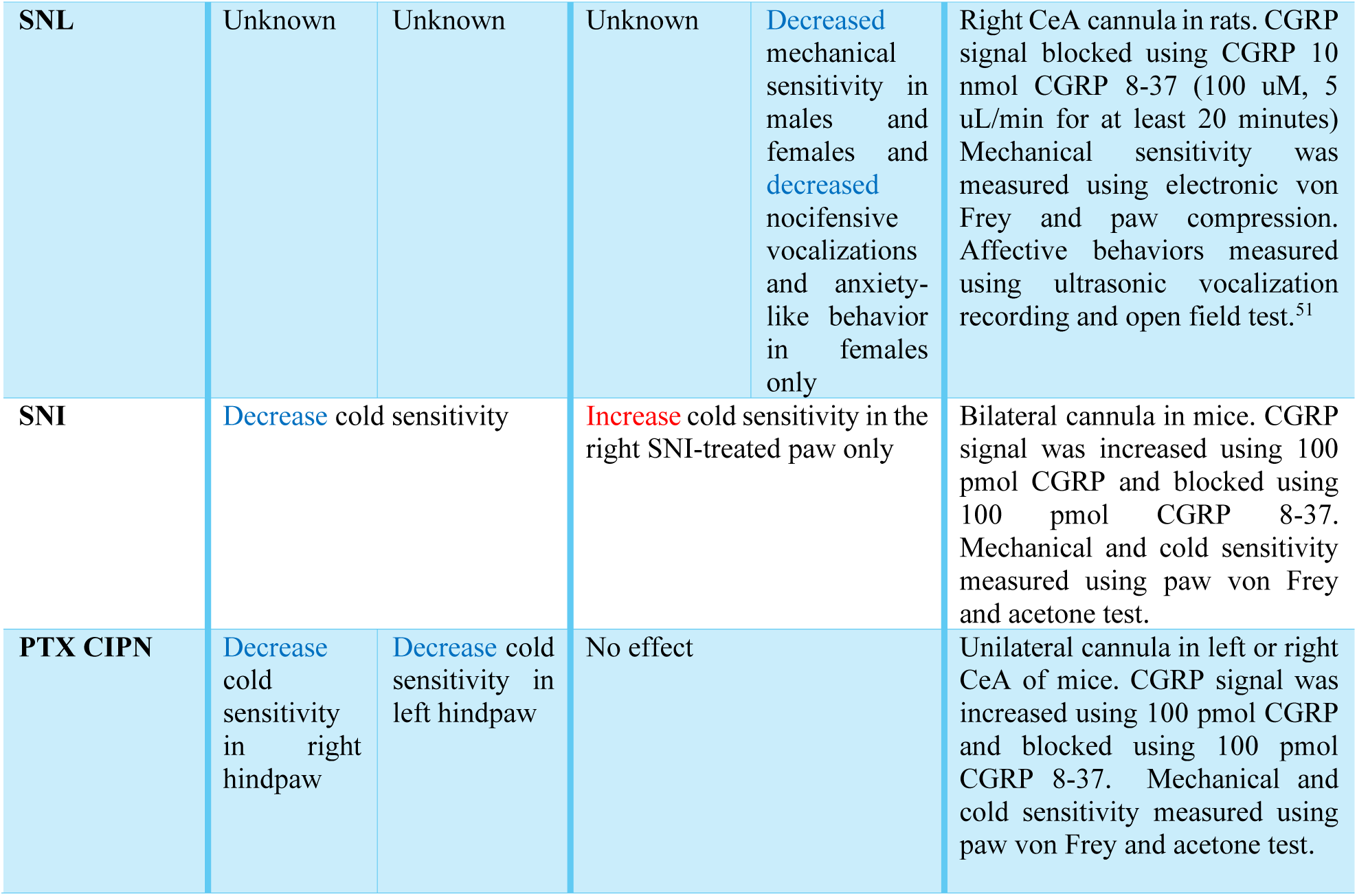
Summary of Known Amygdalar CGRP Effects.

Our study’s data differs from the results of previous publications manipulating CGRP signaling in naïve animals^47,49^ (**Table 2**). A possible explanation for this discrepancy would be the variance of the drug dosage and species tested. Across multiple studies, the dosages of intra-amygdalar CGRP varied from 0.2-2 nmol in rats^49^, 7.5-10 nmol via microdialysis probe in rats^48,51^, 100 pmol in mice^31^, 288 pmol in mice^30,46^, and 750 pmol via microdialysis probe in rats^47^. Although the CGRP/CGRP 8-37 dosage for our investigation was on the low end of published literature (100 pmol), this may be beneficial with regard to specificity. With higher drug dosages injected directly into the brain, there is a possibility that the phenotype observed is caused by non-specific secondary effects rather than on-target effects, including leakage of the CGRP outside of the CeA. Thus, there is more confidence that the observed phenotypes here were a result of CGRP-amygdalar interactions rather than off-target effects. A caveat to this, however, is that a smaller drug dosage of a receptor antagonist could leave some receptor signaling intact.

Based on these results, it appears that amygdalar CGRP has a strong association with modulation of cold sensitivity. In naïve mice, there was a small reduction in cold sensitivity in response to CGRP infusion into the CeA, but a larger reduction was seen in both the SNI and PTX-treated mice, especially in the paw contralateral to the manipulated CeA. This difference in the efficacy of exogenous CGRP may be attributed to a change in CGRP expression between the naïve and neuropathic pain-like states. It is possible that CGRP expression in the PBN to CeA pathway is downregulated during neuropathic pain-like states while CGRP receptors are regulated similarly to the naïve mice, resulting in the potential for a greater anti-nociceptive effect in response to an increase of intra-amygdalar CGRP concentrations. This is not a novel concept in the context of chronic pain-like states, as previous studies have found that mice given CYP to induce bladder pain-like symptoms exhibited an overall reduction in CGRP expression within the bilateral CeA, with a higher reduction present within the left CeA^31^. However, another study found an increase in CGRP mRNA expression in the right CeA of male rats in the acute phase of the SNL model of neuropathic pain and an increase of CGRP and CGRP receptor component mRNA expression in female SNL rats^51^. Immunohistological examination would need to be conducted in the future to confirm this hypothesis and determine if amygdalar CGRP expression in the PTX and SNI models is distinct from the changes seen in the SNL model.

Another interesting result is the difference in behavioral response to amygdalar CGRP injection between the two neuropathic pain models. The most obvious difference is in the responsiveness to CGRP 8-37. The SNI-treated mice exhibited an increase in cold sensitivity in response to CGRP 8-37 in the right SNI-treated hindpaw only, while CGRP 8-37 had no effect on PTX-treated mice. This difference may also be attributed to a difference in CGRP expression within the PBN to CeA pathway, as a downregulation of CGRP may result in a minimal functional effect in CGRP receptor antagonist administration. In other words, if the CeA does not have endogenous peptide signaling, peptide antagonist administration will not produce a functional effect because there are no available ligands to be blocked. Thus, we hypothesize that PTX treatment has the potential to decrease CGRP expression to a greater magnitude than SNI treatment. This difference could be possible given the difference in etiology between PTX and SNI-induced neuropathy. SNI is a focal lesion, causing neuropathy in a single hindlimb (i.e., mononeuropathy), so it may be less likely to cause major effects to other areas of the peripheral nervous system (PNS) or the CNS. On the other hand, PTX is a systemically administered drug, so it is possible that PTX may have an effect outside of the peripheral neuropathy-like symptoms previously reported. Research is currently divided as to whether PTX is able to cross the blood-brain barrier (BBB) and thus cause neurotoxic changes within the CNS^66–68^. Additional experiments need to be conducted to determine if PTX treatment can modify CGRP expression and/or cross the BBB after intraperitoneal administration.

Although male and female mice were used in this study, data were not separately analyzed by sex. Previous studies suggests that CGRP regulation within the PNS or CNS is altered by sex hormones in a complex manner^69–72^. A previous study found an increase in CGRP and CGRP1 receptor components (calcitonin-like receptor (CLR) and receptor activity modifying protein 1 (RAMP1)) mRNA expression in chronic SNL female rats only^51^. Given those results, it is imperative that any future study evaluating the effects of CGRP power to detect potential sex differences.

It is interesting that there was not a strong lateralization effect of CGRP within the amygdala, as there are other studies indicating that neuropathic pain may be modulated in a lateralized manner. For example, one study found that the left CeA was dominantly active during acute neuropathy, but the right CeA was active during chronic-like neuropathy^73^. On the other hand, another study found that the CeA functioned in a more contralateral/ipsilateral manner, similar to our study’s PTX findings^24^. Specifically, lidocaine-mediated right CeA inactivation reduced mechanical and cold sensitivity of the left SNI-treated hindpaw, left and/or right CeA inactivation reduced mechanical sensitivity of the right SNI-treated hindpaw, and inactivation of the left and/or right CeA had no effect on cold sensitivity of the right SNI-treated hindpaw.^24^

CGRP is a promising target to understand the mechanisms behind neuropathic pain, specifically neuropathy-induced cold allodynia. It is important to note that evaluating two models of neuropathic pain with different etiology resulted in different behavioral phenotypes in response to amygdalar CGRP manipulation. Overall, this study indicates that the relationship between CGRP and pain processing within the CeA is complex and possibly dependent on multiple factors, including the subtype of pain (nociceptive, neuropathic, nociplastic), cause of the pain (injury, drug, maladaptive plasticity, etc.), and hemispheric orientation of the CeA.

## ATTRIBUTIONS

The naïve and PTX experiments were conducted by A Trail and J Widner with experimental support provided by L Lewter and S Tack. The SNI data was collected by H Allen and T Nelson. Statistical analysis and figure preparation for all experiments was conducted by A Trail and H Allen. A Trail, H Allen, J Widner, and B Kolber designed experiments. A Trail, H Allen, and B Kolber wrote the manuscript. All authors edited the manuscript.

## ACKNOWLEDGEMENTS

We would like to thank Bradley Taylor for providing C57BL/6J mice for the SNI experiment. Figure illustrations were created in BioRender.

