## Supplemental Table 1 for "Amygdalar Calcitonin Gene-Related Peptide Driven Effects of Cold Sensitivity Induced by Peripheral Neuropathy in Mice"

| Supplemental Table 1. Summary of statistical tests for all figures. | | | | | |
|---|---|---|---|---|---|
| Figure | Comparison | Statistical test(s) | Main Effects | Multiple Comparisons | N |
| 1B | Left paw mechanical sensitivity pre- and post-drug injection into the left CeA | Repeated measures two-way ANOVA with Šidák's multiple comparison test | Time x Treatment p=0.9739<br>Time p=0.0286<br>Treatment p=0.2540<br>Subject p=0.2271 | BL-Post-drug<br>aCSF p=0.4190<br>CGRP p=0.6120<br>CGRP 8-37 p=0.4342 | N=18 (6 female, 12 male) |
| 1C | Right paw mechanical sensitivity pre- and post-drug injection into the left CeA | Repeated measures two-way ANOVA with Šidák's multiple comparison test | Time x Treatment p=0.1241<br>Time p=0.0198<br>Treatment p=0.8142<br>Subject p=0.3048 | BL-Post-drug<br>aCSF p=0.9784<br>CGRP p=0.8498<br>CGRP 8-37 p=0.0100 | N=18 (6 female, 12 male) |
| 1D | Left paw cold sensitivity pre- and post-drug injection into the left CeA | Repeated measures two-way ANOVA with Šidák's multiple comparison test | Time x Treatment p=0.8459<br>Time p=0.0051<br>Treatment p=0.4711<br>Subject p<0.0001 | BL-Post-drug<br>aCSF p=0.5399<br>CGRP p=0.1869<br>CGRP 8-37 p=0.1526 | N=18 (6 female, 12 male) |
| 1E | Right paw cold sensitivity pre- and post-drug | Repeated measures two-way ANOVA with Šidák's | Time x Treatment p=0.1661 | BL-Post-drug<br>aCSF p=0.4109 | N=18 (6 female, 12 male) |

| | | | | | |
|---|---|---|---|---|---|
| | injection into the left CeA | multiple comparison test | Time<br><br>p=0.7949<br><br>Treatment<br><br>p=0.1872<br><br>Subject<br><br>p=0.5013 | CGRP p=0.6667<br><br>CGRP 8-37 p=0.7892 | |
| 1F | Left paw mechanical sensitivity pre- and post-drug injection into the right CeA | Repeated measures two-way ANOVA with Šidák's multiple comparison test | Time x Treatment<br><br>p=0.8333<br><br>Time<br><br>p=0.1815<br><br>Treatment<br><br>p=0.3302<br><br>Subject<br><br>p=0.0587 | BL-Post-drug<br><br>aCSF p=0.9523<br><br>CGRP p=0.5068<br><br>CGRP 8-37 p=0.9081 | N=16 (5 female, 11 male) |
| 1G | Right paw mechanical sensitivity pre- and post-drug injection into the right CeA | Repeated measures two-way ANOVA with Šidák's multiple comparison test | Time x Treatment<br><br>p=0.8117<br><br>Time<br><br>p=0.1086<br><br>Treatment<br><br>p=0.0623<br><br>Subject<br><br>p=0.2522 | BL-Post-drug<br><br>aCSF p=0.9597<br><br>CGRP p=0.6619<br><br>CGRP 8-37 p=0.4579 | N=16 (5 female, 11 male) |
| 1H | Left paw cold sensitivity pre- and post-drug injection into the right CeA | Repeated measures two-way ANOVA with Šidák's multiple comparison test | Time x Treatment<br><br>p=0.8321<br><br>Time<br><br>p=0.2371<br><br>Treatment<br><br>p=0.1118 | BL-Post-drug<br><br>aCSF p=0.9947<br><br>CGRP p=0.6472<br><br>CGRP 8-37 p=0.8177 | N=16 (5 female, 11 male) |

| | | | | | |
|---|---|---|---|---|---|
| | | | Subject<br><br>p=0.0038 | | |
| 1I | Right paw cold sensitivity pre- and post-drug injection into the right CeA | Repeated measures two-way ANOVA with Šidák's multiple comparison test | Time x Treatment<br><br>p=0.2427<br><br>Time<br><br>p=0.0247<br><br>Treatment<br><br>p=0.0362<br><br>Subject<br><br>p=0.0131 | BL-Post-drug<br><br>aCSF p=0.9969<br><br>CGRP p=0.0375<br><br>CGRP 8-37 p=0.5252 | N=16 (5 female, 11 male) |
| 2B | Mechanical sensitivity of the SNI-treated paw pre- and post SNI | Ordinary one-way ANOVA with Dunnett's multiple comparisons test | F=277.7<br><br>p<0.0001 | Pre-SNI vs D14 p<0.0001<br><br>Pre-SNI vs D15 p<0.0001<br><br>Pre-SNI vs D16 p<0.0001<br><br>Pre-SNI vs D17 p<0.0001<br><br>Pre-SNI vs D18 p<0.0001<br><br>Pre-SNI vs D19 p<0.0001 | N=32 (16 left SNI (8 female, 8 male), 16 right SNI (8 female, 8 male) |
| 2C | Cold sensitivity of the SNI-treated paw pre- and post SNI | Ordinary one-way ANOVA with Dunnett's multiple comparisons test | F=106.4<br><br>p<0.0001 | Pre-SNI vs D14 p<0.0001<br><br>Pre-SNI vs D15 p<0.0001<br><br>Pre-SNI vs D16 p<0.0001<br><br>Pre-SNI vs D17 p<0.0001<br><br>Pre-SNI vs D18 p<0.0001<br><br>Pre-SNI vs D19 p<0.0001 | N=32 (16 left SNI (8 female, 8 male), 16 right SNI (8 female, 8 male) |

| | | | | | |
|---|---|---|---|---|---|
| 2D | Left SNI paw mechanical sensitivity pre- and post-drug injection into the left CeA | Repeated measures two-way ANOVA with Šidák's multiple comparison test | Time x Treatment<br><br>p=0.1243<br><br>Time<br><br>p=0.5784<br><br>Treatment<br><br>p=0.0130<br><br>Subject<br><br>p=0.0583 | BL-Post-drug<br><br>aCSF p=0.2661<br><br>CGRP p=0.9370<br><br>CGRP 8-37 p=0.5244 | N=16 (8 female, 8 male) |
| 2E | Right SNI paw mechanical sensitivity pre- and post-drug injection into the left CeA | Repeated measures two-way ANOVA with Šidák's multiple comparison test | Time x Treatment<br><br>p=0.5607<br><br>Time<br><br>p=0.3129<br><br>Treatment<br><br>p=0.6480<br><br>Subject<br><br>p=0.1458 | BL-Post-drug<br><br>aCSF p=0.9899<br><br>CGRP p=0.7710<br><br>CGRP 8-37 p=0.5930 | N=16 (8 female, 8 male) |
| 2F | Left SNI paw cold sensitivity pre- and post-drug injection into the left CeA | Repeated measures two-way ANOVA with Šidák's multiple comparison test | Time x Treatment<br><br>$p<0.0001$<br><br>Time<br><br>p=0.0125<br><br>Treatment<br><br>p=0.0029<br><br>Subject<br><br>p=0.0053 | BL-Post-drug<br><br>aCSF p=0.8612<br><br>CGRP $p<0.0001$<br><br>CGRP 8-37 p=0.1814 | N=16 (8 female, 8 male) |
| 2G | Right SNI paw cold sensitivity pre- and post-drug injection into the left CeA | Repeated measures two-way ANOVA with Šidák's multiple comparison test | Time x Treatment<br><br>$p<0.0001$<br><br>Time | BL-Post-drug<br><br>aCSF p=0.9993<br><br>CGRP $p<0.0001$ | N=16 (8 female, 8 male) |

| | | | | | |
|---|---|---|---|---|---|
| | | | p=0.0002<br>Treatment<br>p<0.0001<br>Subject<br>p=0.0002 | CGRP 8-37 p=0.0129 | |
| 2H | Left SNI paw mechanical sensitivity pre- and post-drug injection into the right CeA | Repeated measures two-way ANOVA with Šidák's multiple comparison test | Time x Treatment<br>p=0.5353<br>Time<br>p=0.0065<br>Treatment<br>p=0.1954<br>Subject<br>p=0.1977 | BL-Post-drug<br>aCSF p=0.1562<br>CGRP p=0.0693<br>CGRP 8-37 p=0.8619 | N=16 (8 female, 8 male) |
| 2I | Right SNI paw mechanical sensitivity pre- and post-drug injection into the right CeA | Repeated measures two-way ANOVA with Šidák's multiple comparison test | Time x Treatment<br>p=0.5612<br>Time<br>p=0.8397<br>Treatment<br>p=0.7727<br>Subject<br>p=0.0002 | BL-Post-drug<br>aCSF p=0.8328<br>CGRP p=0.8919<br>CGRP 8-37 p=0.9573 | N=16 (8 female, 8 male) |
| 2J | Left SNI paw cold sensitivity pre- and post-drug injection into the right CeA | Repeated measures two-way ANOVA with Šidák's multiple comparison test | Time x Treatment<br>p<0.0001<br>Time<br>p=0.0008<br>Treatment<br>p=0.0004<br>Subject | BL-Post-drug<br>aCSF p=0.9315<br>CGRP p<0.0001<br>CGRP 8-37 p=0.4245 | N=16 (8 female, 8 male) |

| | | | | | |
|---|---|---|---|---|---|
| | | | p=0.0009 | | |
| 2K | Right SNI paw cold sensitivity pre- and post-drug injection into the right CeA | Repeated measures two-way ANOVA with Šidák's multiple comparison test | Time x Treatment p<0.0001<br>Time p=0.2788<br>Treatment p=0.0661<br>Subject p=0.0032 | BL-Post-drug<br>aCSF p=0.9989<br>CGRP p<0.0001<br>CGRP 8-37 p=0.0004 | N=16 (8 female, 8 male) |
| 3B | Average mechanical sensitivity pre- and post PTX treatment | Ordinary one-way ANOVA with Dunnett's multiple comparisons test | F=25.76 p<0.0001 | Pre-PTX vs D8 p<0.0001<br>Pre-PTX vs D10 p<0.0001<br>Pre-PTX vs D12 p<0.0001 | N=23 (12 female, 11 male) |
| 3C | Average cold sensitivity pre- and post PTX treatment | Ordinary one-way ANOVA with Dunnett's multiple comparisons test | F=12.82 p<0.0001 | Pre-PTX vs D8 p<0.0027<br>Pre-PTX vs D10 p<0.0001<br>Pre-PTX vs D12 p<0.0001 | N=23 (12 female, 11 male) |
| 3D | Left paw mechanical sensitivity pre- and post-drug injection into the left CeA post-PTX treatment | Repeated measures two-way ANOVA with Šidák's multiple comparison test | Time x Treatment p=0.7573<br>Time p=0.2192<br>Treatment p=0.8092<br>Subject p=0.1292 | BL-Post-drug<br>aCSF p=0.6375<br>CGRP p=0.9992<br>CGRP 8-37 p=0.7100 | N=12 (6 female, 6 male) |
| 3E | Right paw mechanical sensitivity pre- and post-drug injection into the | Repeated measures two-way ANOVA with Šidák's | Time x Treatment p=0.8989 | BL-Post-drug<br>aCSF p=0.9617 | N=12 (6 female, 6 male) |

| | | | | | |
|---|---|---|---|---|---|
| | left CeA post-PTX treatment | multiple comparison test | Time p=0.9019<br>Treatment p=0.9229<br>Subject p=0.5147 | CGRP p>0.9999<br>CGRP 8-37 p=0.9969 | |
| 3F | Left paw cold sensitivity pre- and post-drug injection into the left CeA post-PTX treatment | Repeated measures two-way ANOVA with Šidák's multiple comparison test | Time x Treatment p=0.9265<br>Time p=0.9923<br>Treatment p=0.0007<br>Subject p=0.1447 | BL-Post-drug<br>aCSF p=0.9887<br>CGRP p>0.9999<br>CGRP 8-37 p=0.9913 | N=12 (6 female, 6 male) |
| 3G | Right paw cold sensitivity pre- and post-drug injection into the left CeA post-PTX treatment | Repeated measures two-way ANOVA with Šidák's multiple comparison test | Time x Treatment p=0.3820<br>Time p=0.0021<br>Treatment p=0.5685<br>Subject p=0.1827 | BL-Post-drug<br>aCSF p=0.4050<br>CGRP p=0.0128<br>CGRP 8-37 p=0.5195 | N=12 (6 female, 6 male) |
| 3H | Left paw mechanical sensitivity pre- and post-drug injection into the right CeA post-PTX treatment | Repeated measures two-way ANOVA with Šidák's multiple comparison test | Time x Treatment p=0.2712<br>Time p=0.2781<br>Treatment p=0.9927 | BL-Post-drug<br>aCSF p>0.9999<br>CGRP p>0.9999<br>CGRP 8-37 p=0.1594 | N=11 (6 female, 5 male) |

| | | | | | |
|---|---|---|---|---|---|
| | | | Subject<br><br>p=0.3064 | | |
| 3I | Right paw mechanical sensitivity pre- and post-drug injection into the right CeA post-PTX treatment | Repeated measures two-way ANOVA with Šidák's multiple comparison test | Time x Treatment<br><br>p=0.5838<br><br>Time<br><br>p=0.2957<br><br>Treatment<br><br>p=0.2971<br><br>Subject<br><br>p=0.1385 | BL-Post-drug<br><br>aCSF p=0.8454<br><br>CGRP p=0.9971<br><br>CGRP 8-37 p=0.5063 | N=11 (6 female, 5 male) |
| 3J | Left paw cold sensitivity pre- and post-drug injection into the right CeA post-PTX treatment | Repeated measures two-way ANOVA with Šidák's multiple comparison test | Time x Treatment<br><br>p=0.1944<br><br>Time<br><br>p=0.0006<br><br>Treatment<br><br>p=0.8793<br><br>Subject<br><br>p=0.4183 | BL-Post-drug<br><br>aCSF p=0.3155<br><br>CGRP p=0.0024<br><br>CGRP 8-37 p=0.4868 | N=11 (6 female, 5 male) |
| 3K | Right paw cold sensitivity pre- and post-drug injection into the right CeA post-PTX treatment | Repeated measures two-way ANOVA with Šidák's multiple comparison test | Time x Treatment<br><br>p=0.9356<br><br>Time<br><br>p=0.0102<br><br>Treatment<br><br>p=0.8587<br><br>Subject<br><br>p=0.1802 | BL-Post-drug<br><br>aCSF p=0.4688<br><br>CGRP p=0.2047<br><br>CGRP 8-37 p=0.3423 | N=11 (6 female, 5 male) |
| 4A | Cold sensitivity pre- and post-drug injection into the CeA | Repeated measures two-way ANOVA with Šidák's | Time x Treatment | BL-Post-drug<br><br>aCSF p=0.7952 | N=34 (11 female, 23 male) |

| | | | | | |
|---|---|---|---|---|---|
| | ipsilateral to the tested paw of naïve mice | multiple comparison test | p=0.2434<br>Time<br>p=0.0006<br>Treatment<br>p=0.0243<br>Subject<br>p<0.0001 | CGRP p=0.0052<br>CGRP 8-37 p=0.1024 | |
| 4B | Cold sensitivity pre- and post-drug injection into the CeA contralateral to the tested paw of naïve mice | Repeated measures two-way ANOVA with Šidák's multiple comparison test | Time x Treatment<br>p=0.1382<br>Time<br>p=0.4690<br>Treatment<br>p=0.0252<br>Subject<br>p=0.1958 | BL-Post-drug<br>aCSF p=0.5400<br>CGRP p=0.4299<br>CGRP 8-37 p=0.6226 | N=34 (11 female, 23 male) |
| 4C | Cold sensitivity pre- and post-drug injection into the CeA ipsilateral to the SNI-treated paw | Repeated measures two-way ANOVA with Šidák's multiple comparison test | Time x Treatment<br>p<0.0001<br>Time<br>p=0.0141<br>Treatment<br>p=0.0007<br>Subject<br>p<0.0001 | BL-Post-drug<br>aCSF p=0.9262<br>CGRP p<0.0001<br>CGRP 8-37 p<0.0001 | N=32 (16 left SNI (8 female, 8 male), 16 right SNI (8 female, 8 male) |
| 4D | Cold sensitivity pre- and post-drug injection into the CeA contralateral to the SNI-treated paw | Repeated measures two-way ANOVA with Šidák's multiple comparison test | Time x Treatment<br>p<0.0001<br>Time<br>p<0.0001<br>Treatment | BL-Post-drug<br>aCSF p=0.9564<br>CGRP p<0.0001<br>CGRP 8-37 p=0.0070 | N=32 (16 left SNI (8 female, 8 male), 16 right SNI (8 female, 8 male) |

| | | | | | |
|---|---|---|---|---|---|
| | | | p<0.0001<br>Subject<br>p<0.0001 | | |
| 4E | Cold sensitivity pre- and post-drug injection into the CeA ipsilateral to the tested paw of PTX-treated mice | Repeated measures two-way ANOVA with Šidák's multiple comparison test | Time x Treatment<br>p=0.6420<br>Time<br>p=0.0136<br>Treatment<br>p=0.3202<br>Subject<br>p=0.0292 | BL-Post-drug<br>aCSF p=0.7789<br>CGRP p=0.0956<br>CGRP 8-37 p=0.4482 | N=23 (12 female, 11 male) |
| 4F | Cold sensitivity pre- and post-drug injection into the CeA contralateral to the tested paw of PTX-treated mice | Repeated measures two-way ANOVA with Šidák's multiple comparison test | Time x Treatment<br>p=0.0689<br>Time<br>p<0.0001<br>Treatment<br>p=0.5524<br>Subject<br>p=0.2158 | BL-Post-drug<br>aCSF p=0.0924<br>CGRP p<0.0001<br>CGRP 8-37 p=0.1932 | N=23 (12 female, 11 male) |
